# Alternative Polyadenylation Releases PCBP1-Mediated Suppression of CFIm25 During Macrophage Differentiation

**DOI:** 10.1101/2025.06.19.660645

**Authors:** María del Pilar Mendoza Martín, Salwa Mohd Mostafa, Atish Barua, Claire L. Moore, Srimoyee Mukherjee

## Abstract

CFIm25, a key component of the CFIm complex needed for mRNA 3’ end processing, shows increased protein expression during monocyte-to-macrophage differentiation despite stable mRNA levels. We demonstrate that PCBP1 suppresses CFIm25 translation in monocytes by binding to its long 3’UTR. During differentiation, alternative polyadenylation generates a shorter CFIm25 3’UTR lacking PCBP1 binding sites. RNA immunoprecipitation confirms PCBP1 binding to the long 3’UTR, while ribosome association analysis shows enhanced translation without this interaction. PCBP1 knockdown increases CFIm25 protein specifically in undifferentiated cells and induces macrophage differentiation markers without stimulation. These findings reveal how alternative polyadenylation controls CFIm25 expression during immune cell differentiation by modulating RNA-binding protein interactions and provide insight into post-transcriptional regulation of RNA processing factors.

## Introduction

The cleavage factor Im (CFIm) complex represents a central regulatory node in the cellular RNA processing machinery that controls the choice of poly(A) sites through a process called alternative polyadenylation (APA). CFIm25 (NUDT21/CPSF5), the essential 25-kDa subunit of this complex, recognizes UGUA sequence motifs upstream of polyadenylation sites and enhances recruitment of the polyadenylation machinery (1,2). While APA impacts over 70% of human genes (3), CFIm25 specifically regulates a substantial subset of these events, influencing mRNA stability, localization, and translation efficiency. Monocyte-to-macrophage differentiation, a process critical for innate immune function, is accompanied by widespread alterations in polyadenylation patterns (4,5). Notably, CFIm25 protein levels increase substantially during this transition (4), but the mechanisms governing CFIm25 expression itself remain poorly characterized.

In general, the precise regulation of all of the C/P factors themselves remains poorly understood, despite their critical importance in gene expression control. These factors are regulated at multiple levels (6-8). For example, changes in levels of each component alter the protein complex stoichiometry and shift poly(A) site choice (6). These changes can be mediated transcriptionally, as seen with CstF64 upregulation during B-cell activation and differentiation (9) or by ubiquitination and degradation of Pcf11 or CPSF73 to drive APA and regulate cancer cell properties (10,11). Other post-translational modifications like sumoylation and the cell cycle-dependent phosphorylation of the poly(A) polymerase (10,12), and ubiquitination of Pcf11 and CPSF73 by cancer-specific ubiquitin ligases that drive APA and affect tumor progression (11,13); and through protein complex stoichiometry, where changes in levels of each component can shift poly(A) site choice (6). Ubiquitination and lactylation of CFIm25 reprograms APA in renal fibrosis and cancer respectively (14,15). In some cases, such as with PCF11 and CstF77, the use of intronic poly(A) sites can alter the amount of full-length, functional protein (16-18). However, translational regulation of C/P factors remains one of the least explored mechanisms. The action of these regulators can be blocked by APA that removes their binding sites in the 3’ UTR. A recent study by Khan et al. demonstrated that CFIm25 undergoes APA to generate multiple mRNA isoforms differentially regulated by miRNAs, providing evidence for translational control of this key C/P factor (19). However, regulation of CFIm25 translation by RNA-binding proteins (RBPs) has not been addressed.

RBPs act through sequence-specific interactions with mRNA untranslated regions (20). These regulatory factors can fine-tune protein output through varied mechanisms including mRNA stability control and repression or enhancement of translation. PCBP1 (Poly(C)-binding protein 1) represents a well-characterized translational regulator that binds C-rich and GC-rich motifs within mRNA untranslated regions (21). This hnRNP E family member has been implicated in both mRNA stabilization and translational suppression across diverse cellular contexts. PCBP1’s regulatory activity typically occurs without requiring changes in its own expression levels during cellular transitions, making it an ideal candidate for context-dependent regulation of translation.

Here we investigate whether PCBP1 regulates CFIm25 expression through interaction with its 3’UTR. CFIm25 presents a particularly interesting case study, as it not only undergoes robust protein upregulation during macrophage differentiation despite stable mRNA levels but also represents a core component of the machinery that determines 3’UTR length through alternative polyadenylation. Our findings demonstrate that PCBP1 binding serves as a mechanism to regulate CFIm25 protein levels during differentiation. Importantly, this molecular circuit directly impacts monocyte differentiation, with PCBP1 functioning as a regulator of differentiation at least in part through its effects on CFIm25 translation. This study establishes a molecular link between post-transcriptional regulation of CFIm25 and immune cell differentiation.

## Materials and Methods

### Cell Culture and Differentiation

THP-1 cells were maintained in RPMI 1640 medium supplemented with 10% heat-inactivated fetal bovine serum, 2 mM L-glutamine, and antibiotics at 37°C in a humidified environment with 5% CO_2_. For differentiation, cells were treated with 3 nM phorbol-12-myristate-13-acetate (PMA) for 24 hours. This concentration was selected based on previous optimization studies showing efficient differentiation (4).

### Bioinformatic Analysis of RBP Binding Sites

RBPmap (http://rbpmap.technion.ac.il/) was used to predict RNA-binding protein binding sites with consideration of motif environments. The analysis was performed using the CFIm25 3’UTR sequence with default parameters (stringency: high, conservation filter: on). POSTAR3 (http://postar.ncrnalab.org/) was used to explore experimental evidence for RBP binding from publicly available eCLIP, CLIP-seq, and RNA-seq datasets. Results from both tools were merged to identify high-confidence binding sites present in both analyses. Binding sites were visualized using the UCSC Genome Browser (GRCh38/hg38).

### RNA Analysis and RT-qPCR

Total RNA was isolated using TRIzol reagent following manufacturer’s protocol. For 3’UTR analysis, isoform-specific primers were designed to amplify either the long or total 3’UTR regions of CFIm25. RT-qPCR was performed using SYBR Green with β-actin as internal control. Each sample was analyzed in triplicate, and relative expression was calculated using the 2^-ΔΔCt method. Primer sequences are provided in Supplementary Table 1.

### siRNA Transfection

ON-TARGETplus Human PCBP1 siRNA - SMARTpool or non-targeting control siRNA was transfected into THP-1 cells using Dharmafect following manufacturer’s protocol. Final siRNA pool concentration was 50 nM. Knockdown efficiency was assessed 48 hours post-transfection by both Western blot and RT-qPCR. All experiments were performed with at least three independent transfections.

### RNA Immunoprecipitation

Cells were lysed in RIPA buffer (50 mM Tris-HCl pH 7.4, 150 mM NaCl, 1% NP-40, 0.5% sodium deoxycholate, 0.1% SDS, 1 mM EDTA) supplemented with protease inhibitors and RNase inhibitors. Lysates were immunoprecipitated with antibodies against PCBP1 or RPL26. RNA was purified from immunoprecipitates using TRIzol and analyzed by RT-qPCR. Normal rabbit IgG served as negative control, and input RNA was used for normalization. Antibody details are provided in Supplementary Table 2.

### Western Blot Analysis

Cell lysates were prepared in RIPA buffer supplemented with protease inhibitors. Protein concentration was determined using BCA assay. Equal amounts of protein (50 μg) were resolved by 10% SDS-PAGE and transferred to PVDF membranes. After overnight incubation with primary antibodies at 4°C, membranes were washed and incubated with HRP-conjugated secondary antibodies. Proteins were visualized using enhanced chemiluminescence detection system. Band intensities were quantified using ImageJ software. Antibody details are provided in Supplementary Table 2.

### Differentiation Marker Analysis

Cell attachment was quantified by counting both suspended and adherent cells after gentle washing with PBS. The surface marker expressions were analyzed by flow cytometry using PE-conjugated anti-CD38 and anti-CD11b antibodies (BD Biosciences). Cells were incubated with antibody for overnight at 4°C in the dark, washed twice with PBS containing 2% FBS, analyzed using an Attune flow cytometer (Thermo Fisher Scientific), and data were analyzed using FACS diva. For analysis, samples were gated on light scattering properties to exclude dead cells and debris. Unstained control samples were used to determine the level of background fluorescence. Antibody details are provided in Supplementary Table 2.

## Statistical Analysis

All experiments were performed at least three times independently. Data are presented as mean ± standard deviation. Statistical significance was determined using Student’s t-test. P values < 0.05 were considered statistically significant. Statistical analyses were performed using GraphPad Prism software.

## Results

### CFIm25 Expression is Regulated Through APA and PCBP1 Binding

Our previous studies demonstrated a significant increase in CFIm25 protein levels during monocyte-to-macrophage differentiation across multiple cell systems including U937 and THP-1 cells (5). However, the regulation of CFIm25 transcript abundance during this transition remained unexplored. We examined CFIm25 mRNA levels using both published RNA-seq datasets and RT-qPCR analysis. In contrast to the pronounced protein elevation, CFIm25 mRNA levels showed slight decreases across THP-1, human primary monocytes, and U937 cell lines during differentiation (Supplementary Fig. 1A) (4,5). Analysis of our own 3’ -seq data comparing undifferentiated versus differentiated U937 and HL-60 cells confirmed this pattern, revealing consistent but modest decreases in mRNA levels that cannot account for the large protein increase observed during differentiation (Supplementary Fig. 1B). This discrepancy suggested post-transcriptional regulatory mechanisms.

Upon reviewing our previously published 3’ end sequencing data from U937 cells (5), we observed that CFIm25 undergoes alternative polyadenylation during differentiation (Supplementary Fig. 1C). UCSC genome browser visualization revealed that undifferentiated U937 cells express approximately equal proportions of long and short 3’UTR isoforms (Supplementary Fig. 1D). Following PMA-induced differentiation, there is a clear shift toward utilization of the proximal poly(A) site, resulting in a predominantly shorter 3’UTR.

To validate this observation in our experimental THP-1 model, we performed RT-qPCR analysis using isoform-specific primers (schematic shown in Fig. 1A). These experiments confirmed the significant shift in CFIm25 UTR usage, showing an almost 2-fold decrease in the long-to-short UTR ratio after 24 hours of PMA treatment (Fig. 1B). Importantly, these changes represent genuine alternative polyadenylation rather than differential transcript stability, as total mRNA levels do not increase throughout differentiation.

**Fig 1.**
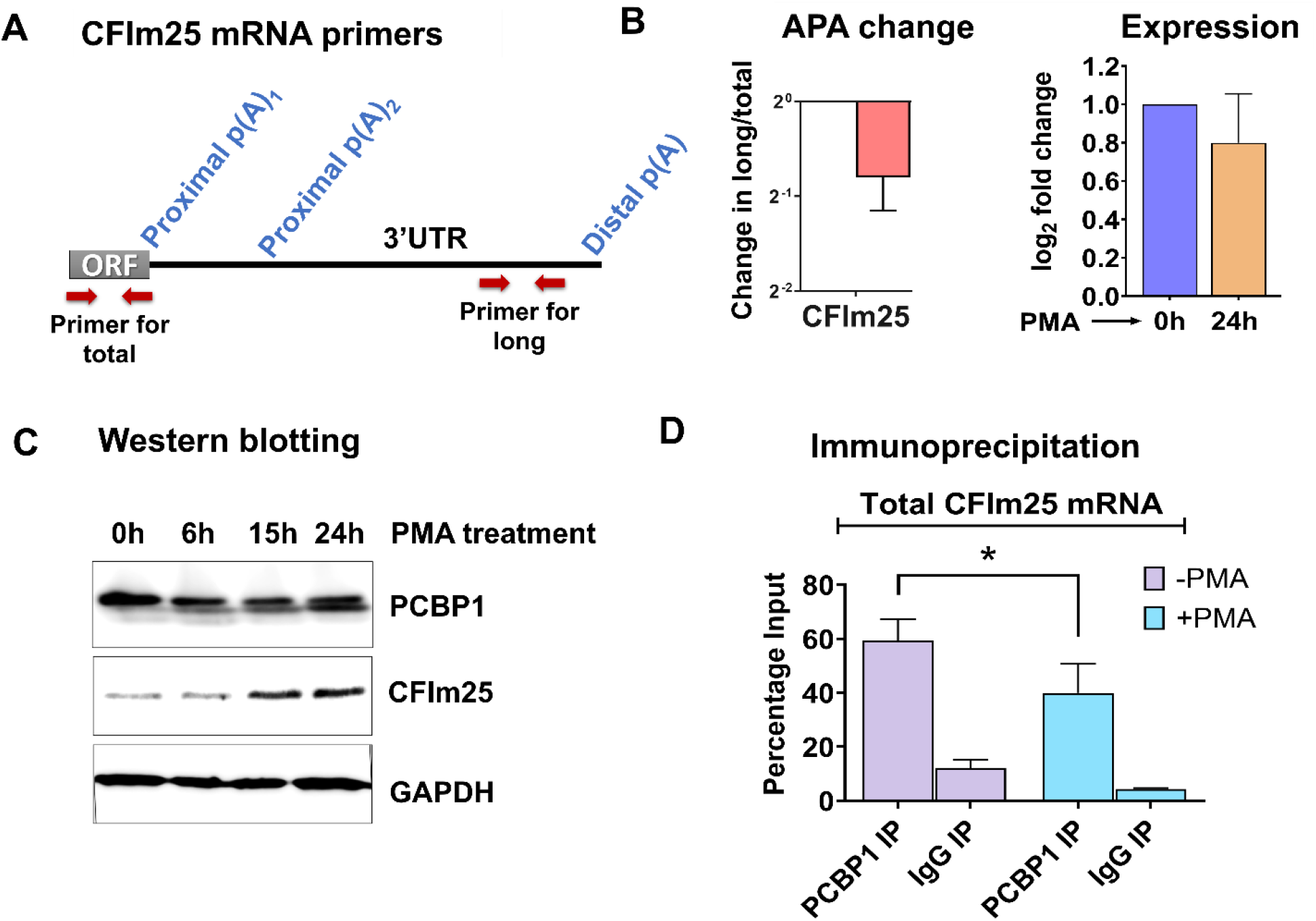
Increased CFIm25 expression correlates with mRNA shortening and reduced PCBP1 binding. (A) The schematic shows the CFIm25 mRNA structure with open reading frame (ORF) and 3’UTR, with two proximal and distal poly(A) sites indicated in blue. Primer pairs for total and long are indicated by red arrows (B) Analysis of CFIm25 in control and PMA-treated THP-1 cells. *Left*: APA analysis showing Log2 ratio of long/total CFIm25 mRNA. *Right*: Log2 fold change in total CFIm25 mRNA levels. Data represent mean ± SD (n=3); p < 0.01. (C) Western blot showing PCBP1 and CFIm25 protein levels during differentiation with PMA treatment at different time points (0h, 6h, 15h, and 24h). GAPDH serves as loading control. (D) RNA immunoprecipitation using PCBP1 or control IgG antibody followed by CFIm25 qPCR for the longest transcript in undifferentiated (-PMA) and differentiated (+PMA) cells. Data normalized to input and represented as percentage input. Statistical significance determined by Student’s t-test.

### PCBP1 Binds Differentially to CFIm25 UTR Isoforms

Given the observed APA shift and the well-established role of RNA-binding proteins (RBPs) in translational regulation through UTR interactions, we investigated whether CFIm25 translation might be controlled through differential RBP binding to its 3’UTR isoforms. We performed comprehensive bioinformatic analysis using RBPmap and POSTAR3 to predict RBP binding sites across the CFIm25 3’UTR. This analysis revealed 92 potential binding sites from RBPmap and 53 from POSTAR3, with 26 high-confidence sites overlapping between both tools (Supplementary Fig. 2A).

Among the predicted RBPs, PCBP1 emerged as a particularly intriguing candidate for several reasons. First, PCBP1 was predicted with high confidence by both tools, suggesting robust binding potential. Second, unlike other candidates that showed distributed binding patterns throughout the 3’UTR, PCBP1 exhibited a well-defined binding site in the distal portion of the UTR that would be specifically lost during APA-mediated shortening (Supplementary Fig. 2B). Third, PCBP1 has an established role in translational repression through UTR binding (22), making it an ideal candidate for investigating APA-mediated translational control.

Western blot analysis confirmed that while CFIm25 levels increase, PCBP1 protein levels remain stable during differentiation (Fig. 1C), supporting the idea that regulatory changes could be due to altered binding site accessibility rather than changes in protein abundance. To validate PCBP1’s interaction with CFIm25 mRNA, we performed RNA immunoprecipitation using a validated PCBP1 antibody. RNA immunoprecipitation demonstrated strong enrichment of CFIm25 transcripts in undifferentiated cells that decreased by more than 20% upon differentiation (Fig. 1D), supporting our hypothesis that PCBP1 binding is modulated by CFIm25 3’ UTR length. To further confirm that PCBP1 specifically associates with the long 3’UTR isoform, we performed RIP followed by qPCR using primers designed just downstream of the PCBP1 binding site in the long CFIm25 isoform. PCBP1 immunoprecipitation showed enrichment for the long isoform specifically, while control IgG showed minimal background binding to either target (Supplementary Fig. 2C). This demonstrates that PCBP1 can bind to CFIm25 transcripts containing the extended 3’UTR region where its binding sites are predicted to reside.

### Functional Consequence of PCBP1 Binding on CFIm25 Translation

To investigate whether PCBP1 mediates translational control of CFIm25, we depleted PCBP1 using siRNA in THP-1 cells. Knockdown efficiency was consistently >80% as assessed by Western blot in both undifferentiated and differentiated cells (Fig. 2A, 2D). In undifferentiated cells, PCBP1 depletion led to an increase in CFIm25 protein levels (Fig. 2A). Importantly, this increase occurred without any significant change in CFIm25 mRNA abundance, consistent with regulation at the translational level (Fig. 2B). The effect was highly specific, as a control siRNA showed no impact on CFIm25 protein levels. In contrast, PCBP1 knockdown had no significant effect on CFIm25 protein or RNA levels in differentiated cells (Fig. 2D-E), consistent with our hypothesis that PCBP1 regulation occurs specifically through binding sites in the long UTR form.

**Fig 2.**
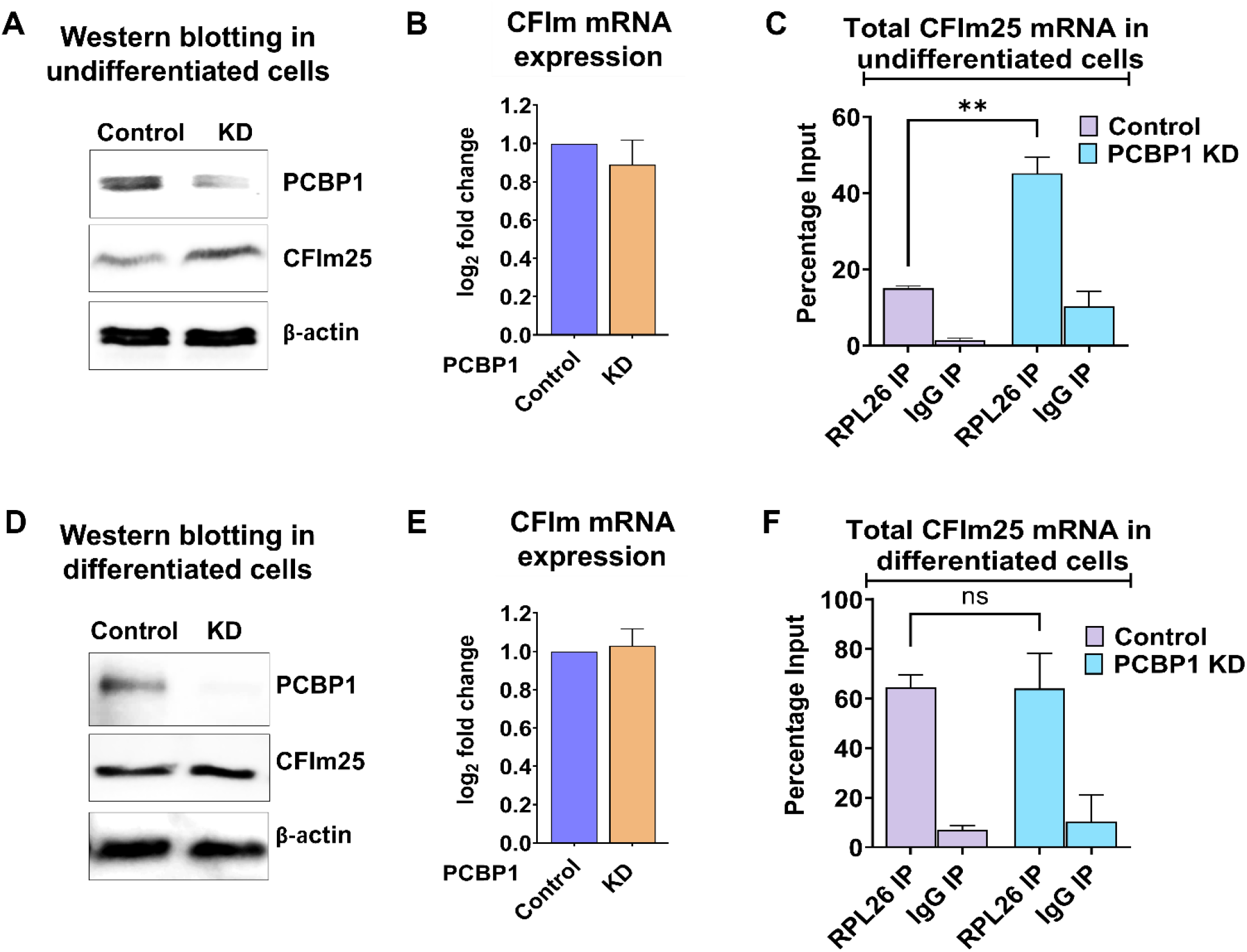
Functional consequences of PCBP1 binding on CFIm25 translation. (A) Western blot analysis of CFIm25 and PCBP1 protein levels in undifferentiated THP-1 cells 48h after transfection with control or PCBP1 siRNA knockdown (KD). PCBP1 KD leads to increased CFIm25 protein levels. β-actin serves as loading control. (B) Log2 fold change of total CFIm25 in control and PCBP1 KD samples at undifferentiated conditions in THP-1 cells. Data represent mean ± SD (n=3) (C) RPL26 RNA immunoprecipitation in undifferentiated cells showing ribosome association of CFIm25 mRNA in control and PCBP1 KD conditions. IgG immunoprecipitation serves as negative control. Data normalized to input and represented as percentage of input. (D) Western blot analysis of CFIm25 and PCBP1 protein levels in differentiated (PMA-treated) THP-1 cells 48h after transfection with control or PCBP1 siRNA KD. PCBP1 knockdown has minimal effect on CFIm25 protein levels in differentiated cells. β-actin serves as loading control. (E) Log2 fold change of total CFIm25 in control and PCBP1 KD samples at differentiated conditions in THP-1 cells. Data represents mean ± SD (n=3) (F) RPL26 RNA immunoprecipitation in differentiated cells showing similar ribosome association of CFIm25 mRNA in both control and PCBP1 KD conditions. IgG immunoprecipitation serves as negative control. Data normalized to input and represented as percentage of input. Statistical significance determined by Student’s t-test.

To assess the impact of PCBP1 on translation, we used RNA immunoprecipitation with antibodies against the ribosomal protein RPL26, a well-validated and reliable marker for ribosome association in human cells (23,24). 18S rRNA served as a positive control and IgG as a negative IP control. In undifferentiated cells, PCBP1 knockdown significantly increased CFIm25 mRNA association with ribosomes compared to control siRNA treatment (Fig. 2C). Control IgG immunoprecipitations showed minimal background binding, confirming the specificity of our RPL26 pulldown. In differentiated cells, however, CFIm25 mRNA showed similar levels of ribosome association regardless of PCBP1 status (Fig. 2F), consistent with the predominance of the shorter UTR isoform lacking PCBP1 binding sites. While we might expect some increase in ribosome association for the remaining long isoforms in PCBP1-depleted differentiated cells, a significant change in their amount may have different outcomes. These results demonstrate that PCBP1 binding to the long 3’UTR suppresses CFIm25 translation by reducing its association with the translational machinery, a mechanism that is specifically lost during differentiation due to APA-mediated removal of binding sites.

Collectively, these results establish that alternative polyadenylation during monocyte differentiation removes PCBP1 binding sites from the CFIm25 3’UTR, releasing translational repression and allowing for increased protein expression without changes in mRNA levels. Our work reveals a distinct mechanism where a key regulator of APA is itself controlled by the process it governs, but through 3’UTR shortening and translational control rather than through intronic polyadenylation.

### PCBP1-CFIm25 Regulatory Axis Controls Monocyte Differentiation

We have previously shown that overexpression of CFIm25 enhanced the efficiency of monocyte differentiation (4), which suggested that the increase in CFIm25 seen upon PCBP1 depletion might have a similar effect. To determine whether the PCBP1-CFIm25 regulatory mechanism functionally impacts monocyte differentiation, we examined whether PCBP1 knockdown alone could induce differentiation-associated phenotypes in unstimulated THP-1 cells. A hallmark of monocyte-to-macrophage differentiation is the transition from suspension growth to adherence as cells develop macrophage characteristics. When quantified, PCBP1 knockdown significantly increased cell attachment (∼20% attached cells) compared to control cells (∼8% attachment), indicating spontaneous differentiation even without PMA treatment (Fig. 3A). Cell viability remained consistent across all conditions (>90%), confirming that the observed effects were not due to cytotoxicity of the siRNA treatments (Fig. 3B).

**Fig 3.**
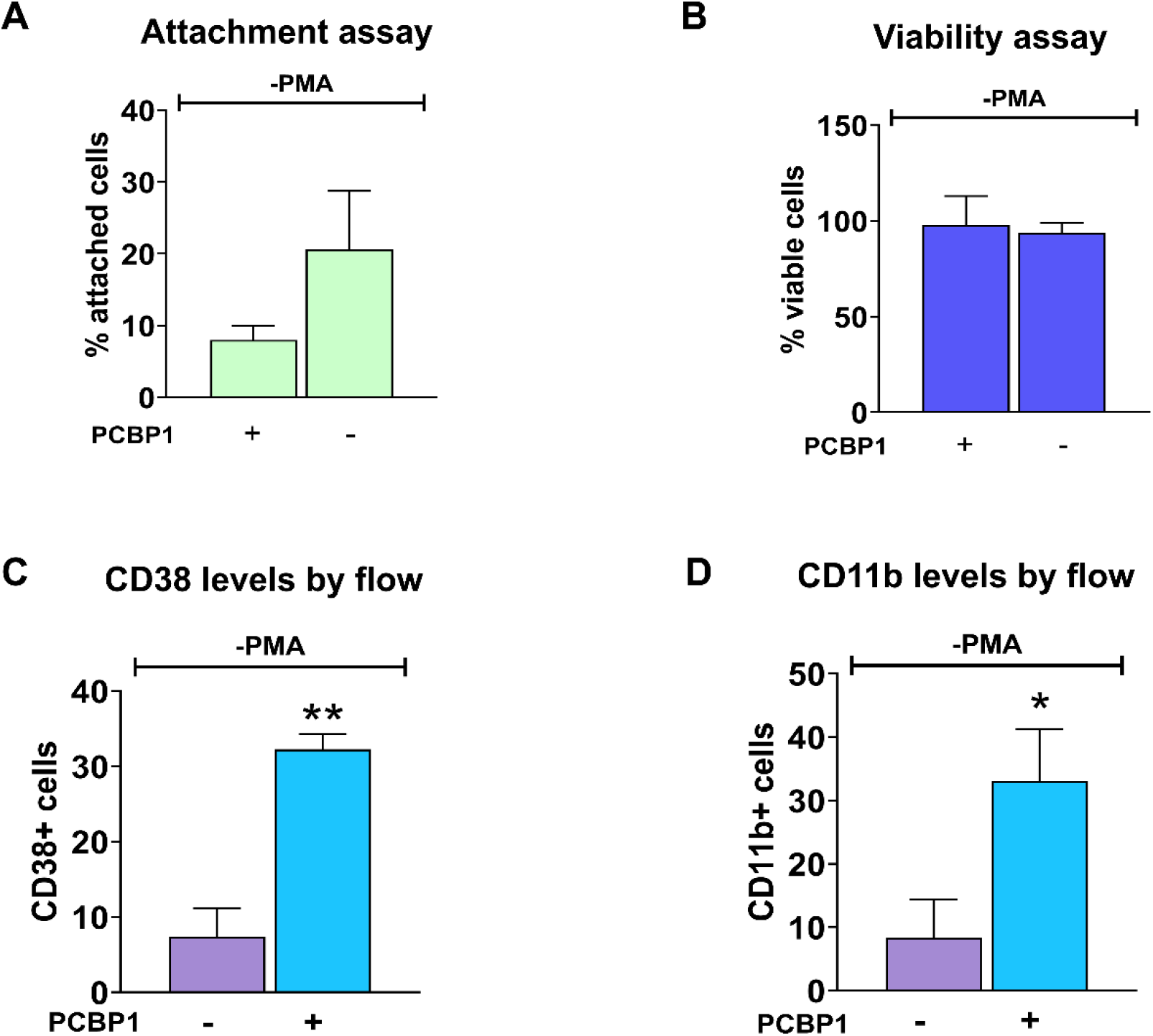
PCBP1-CFIm25 regulatory axis controls monocyte differentiation. THP-1 cells were transfected with control or PCBP1 siRNA and analyzed without PMA treatment. (A) Cell attachment assay quantifying the percentage of adherent cells 48 hours post-transfection. PCBP1 knockdown significantly increases cell adhesion compared to control (mean ± SD, n=3; *p<0.05 by Student’s t-test). (B) Cell viability assay demonstrating that cell death is not affected by PCBP1 depletion. Values represent percentage of viable cells 48 hours post-transfection (mean ± SD, n=3). (C-D) Bar graph summarizing the percentage of CD38+ (C) and CD11b+ (D) cells across three independent experiments, demonstrating significant upregulation of these differentiation markers upon PCBP1 knockdown. Statistical significance determined by Student’s t-test.

To further validate the differentiation phenotype, we measured expression of two well-established macrophage surface markers by flow cytometry. PCBP1 knockdown significantly increased the percentage of CD38-positive cells to approximately 32% compared to 7% in control cells (Fig. 3C, ** p<0.01). Similarly, CD11b expression, another key macrophage differentiation marker, showed a significant increase in PCBP1-depleted cells (33% positive) compared to control cells (8% positive) (Fig. 3D, * p<0.05). These quantitative data from multiple differentiation markers provide strong evidence that PCBP1 depletion induces spontaneous macrophage differentiation in the absence of PMA stimulation.

These results demonstrate that PCBP1 functions as a suppressor of monocyte differentiation, and its depletion is sufficient to induce key differentiation markers. Together with our mechanistic findings showing PCBP1-mediated translational repression of CFIm25, these data establish that the PCBP1-CFIm25 regulatory axis serves as a control mechanism for monocyte-to-macrophage transition.

## Discussion

Our findings reveal a refined regulatory circuit where alternative polyadenylation modulates the translation of CFIm25 by controlling accessibility to PCBP1 binding sites. This mechanism parallels similar regulatory strategies observed in other biological contexts yet represents a novel form of regulation in the context of macrophage differentiation. Most importantly, we demonstrate that this regulatory axis has direct functional consequences for macrophage differentiation, establishing a previously uncharacterized link between post-transcriptional gene regulation in monocytes and cell fate decisions.

The PCBP1-mediated translational control we observed aligns with its well-established regulatory roles across diverse biological contexts. Early work by Wang and colleagues demonstrated that PCBP1 suppresses PRL-3 phosphatase translation through interaction with conserved GCCCAG motifs in the 5’ UTR (22). This regulation occurs independently of mRNA levels, similar to our findings with CFIm25 where PCBP1 appears to inhibit translation without affecting mRNA abundance. Beyond translational control, PCBP1 has diverse functions in RNA processing. Tripathi et al. showed that PCBP1 colocalizes with SMAD3 in nuclear speckles to regulate CD44 alternative splicing during TGF-β-induced epithelial-to-mesenchymal transition (25). Even more relevant to our work, Cheng et al. demonstrated that PCBP1 regulates RNA splicing of STAT3, promoting macrophage-like phenotype modulation of vascular smooth muscle cells (26). Khan et al. showed that miRNAs binding to the 3’ UTR of CFIm25 mRNA decreased its mRNA levels (19). PCBP1 represents another layer of post-transcriptional regulation that can fine-tune expression of critical factors without altering mRNA abundance. These diverse regulatory functions underscore PCBP1’s potential as a master regulator of RNA processing networks rather than a single-target repressor.

The identification of translational control through RBP binding as a regulatory mechanism for CFIm25 significantly expands our understanding of how cleavage/polyadenylation factors are regulated. Transcriptional regulation of these factors has been well-documented - most notably the upregulation of CstF-64 during B-cell activation (9). Translational and post-translational modifications represent another layer of regulation. This mechanism is particularly relevant for rapid cellular transitions, as it obviates the need for transcriptional activation and protein translation from scratch. Collectively, studies, including this one, reveal multiple layers of post-transcriptional control governing CFIm25 expression. These include APA to remove binding sites for inhibitors of mRNA translation and stability, as well as post-translational modifications (cite all of the studies), and highlight the sophistication of regulatory mechanisms controlling C/P factors. While our findings demonstrate that CFIm25 is regulated by alternative polyadenylation, further studies will be needed to determine whether CFIm25 directly influences its own 3’ UTR processing. Regulatory feedback loops have been described for other RNA processing factors. For example, autoregulation of alternative splicing has been observed for SR proteins and hnRNPs (27,28). More directly relevant, Wang et al. recently showed that intronic polyadenylation within the PCF11 controls its own gene expression through an intronic polyadenylation site (16).

From a mechanistic perspective, our data suggests that PCBP1 binding to the 3’UTR interferes with efficient translation of the long CFIm25 isoform, potentially by affecting mRNA localization or interactions with the translation machinery. RBPs bound to 3’UTRs can influence translation through various mechanisms, including modulation of poly(A) tail function, interaction with translation initiation factors, or affecting mRNA localization to translational compartments (29). The fact that differentiated cells show constitutive ribosome association regardless of PCBP1 levels provides compelling evidence that 3’UTR shortening is a key regulatory switch that removes PCBP1-mediated translational repression.

The robust induction of both CD38 and CD11b in PCBP1-depleted cells, along with the morphological transition to adherent growth, provides compelling evidence that this regulatory mechanism is functionally significant for macrophage differentiation. PCBP1 appears to function as a molecular gatekeeper preventing premature differentiation, similar to how SOX2 maintains pluripotency in stem cells by suppressing differentiation genes (30). The demonstration that PCBP1 depletion alone can initiate differentiation without external stimuli suggests that post-transcriptional regulatory networks may play more prominent roles in immune cell fate decisions than previously appreciated. This places PCBP1 among critical developmental checkpoints that must be relieved to enable full cellular transition.

In conclusion, our study reveals a multi-layered regulatory mechanism controlling CFIm25 expression and demonstrates its functional importance in cellular differentiation. This work enhances our understanding of how alternative polyadenylation and RNA-binding proteins can coordinate to regulate key components of the RNA processing machinery, adding a new dimension to our appreciation of post-transcriptional gene regulation networks. Future studies exploring whether similar mechanisms regulate other C/P factors could uncover additional layers of this regulatory complexity and identify new therapeutic targets for conditions involving dysregulated RNA processing.

## Acknowledgments

This work was supported by the National Institutes of Health (NIH) grant 1R01AI152337 (to CM). The authors also acknowledge the Tufts Flow Cytometry Core for technical support.

## Author Contributions

All authors contributed to the study. CM and S Mukherjee conceived the study, while S Mukherjee designed the experiments. CM acquired the funding. Material preparation, data collection and analysis were performed by S Mukherjee and PM. AB helped with the flow cytometry and bioinformatic analysis was performed by S Mostafa. The first draft of the manuscript was written by S Mukherjee and CM provided critical input. All authors commented on previous versions of the manuscript. All authors read and approved of the final manuscript.

