## Supplementary material for "Alternative Polyadenylation Releases PCBP1-Mediated Suppression of CFIm25 During Macrophage Differentiation"

**Supplementary Table 1. Primers used in this study:**

| Primer name | Forward primer 5'-3' | Reverse primer 5'-3' |
|---|---|---|
| CFIm25 total | ACCAGGAGAAGATGAAGTTGA | ATGCACCATTGTTTGAATTGT |
| CFIm25 long | TGGACCCTTGCTTTGATACCA | AGGTCTGTTGCTTGCAAACC |
| ACTB total | CATGTACGTTGCTATCCAGGC | CTCCTTAATGTCACGCACGAT |

**Supplementary Table 2. Antibodies used in this study:**

***For western blot and immunoprecipitation:***

| Catalogue no. | Name | Host | MW | Vendors |
|---|---|---|---|---|
| sc-81109 | CFIm25 | Mouse | 26 kDa | Santa Cruz Biotechnology |
| A1044 | PCBP1 | Rabbit | 37 kDa | Abclonal |
| A300-686A-T | RPL26 | Rabbit | 26 kDa | Bethyl |
| sc-47778 | β-actin | Mouse | 43 kDa | Santa Cruz Biotechnology |
| sc-365062 | GAPDH | Mouse | 37 kDa | Santa Cruz Biotechnology |

***For flow cytometry:***

| Catalogue no. | Name | Host | Fluorophore | Vendor |
|---|---|---|---|---|
| 102707 | CD38 | Mouse | PE | Biolegend |
| 393111 | CD11b | Mouse | PE | Biolegend |

### Supplementary figures:

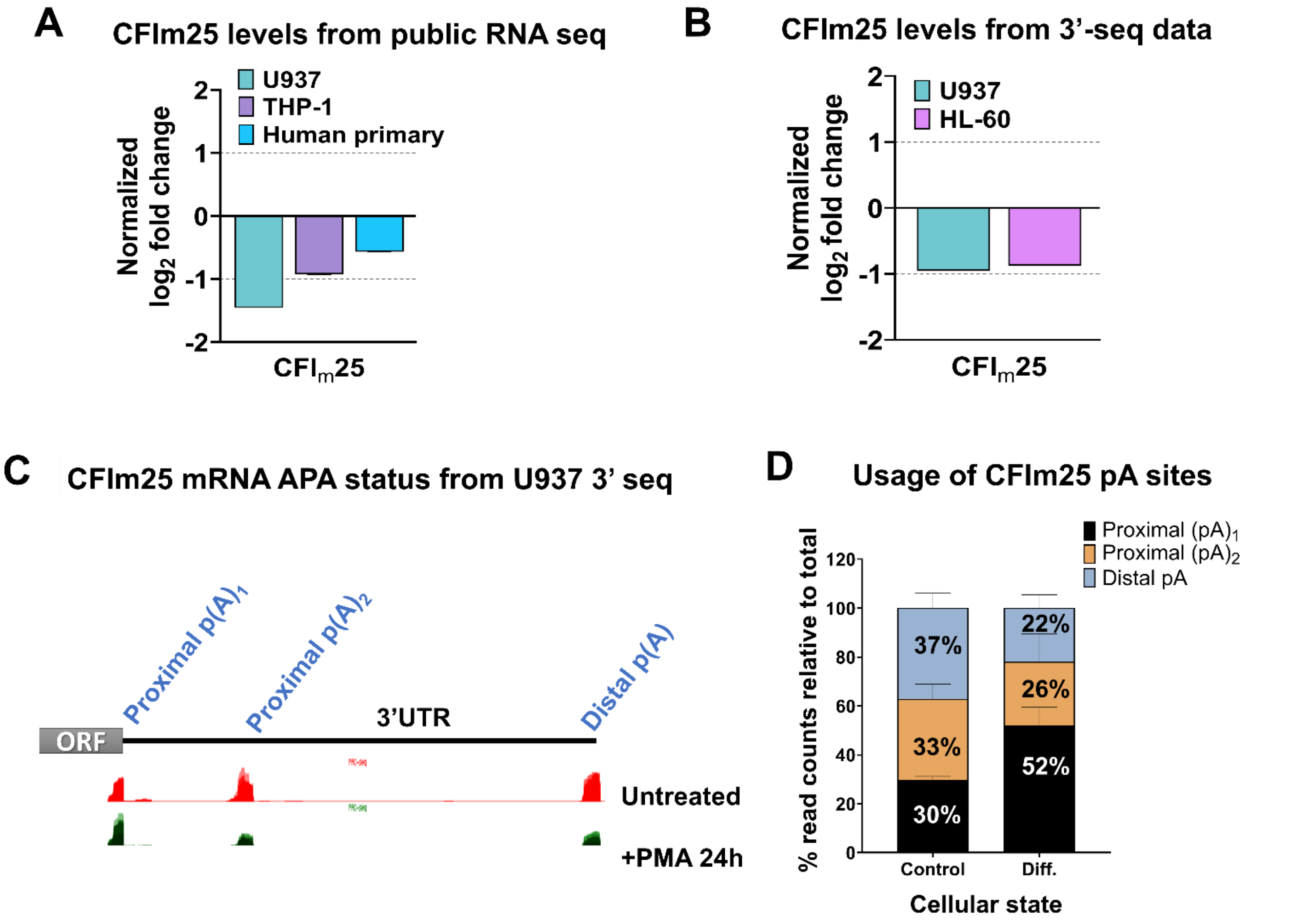


**Supplementary Fig 1. CFIm25 mRNA levels across macrophage cells.** (A) CFIm25 mRNA levels from previously published RNA-seq datasets comparing differentiated versus undifferentiated cells in U937, THP-1 and human primary monocytes. Values represent log2 fold change normalized to undifferentiated controls, showing modest decreases in CFIm25 mRNA levels after differentiation across all cell types. (B) CFIm25 mRNA levels from our RNA-3' end seq data comparing differentiated versus undifferentiated U937 and HL-60 cells. Values represent log2 fold change normalized to undifferentiated controls, confirming consistent but modest decreases in CFIm25 mRNA levels that do not explain the magnitude of protein increases observed during differentiation. (C) UCSC genome browser tracks showing CFIm25 3'UTR usage in undifferentiated (red) and PMA-treated (24h) U937 cells (green). The schematic above shows the CFIm25 mRNA structure with open reading frame (ORF) and 3'UTR, with two proximal and a distal poly(A) site indicated in blue. (D) Quantitative bar graphs reflecting the mean relative usage of CFIm25 poly(A) sites with respect to the total read counts (at all the poly (A) sites) as visualized in the UCSC genome browser. Student's t-test was performed to determine the significance between the treatment groups and P value <0.05 was considered significant.

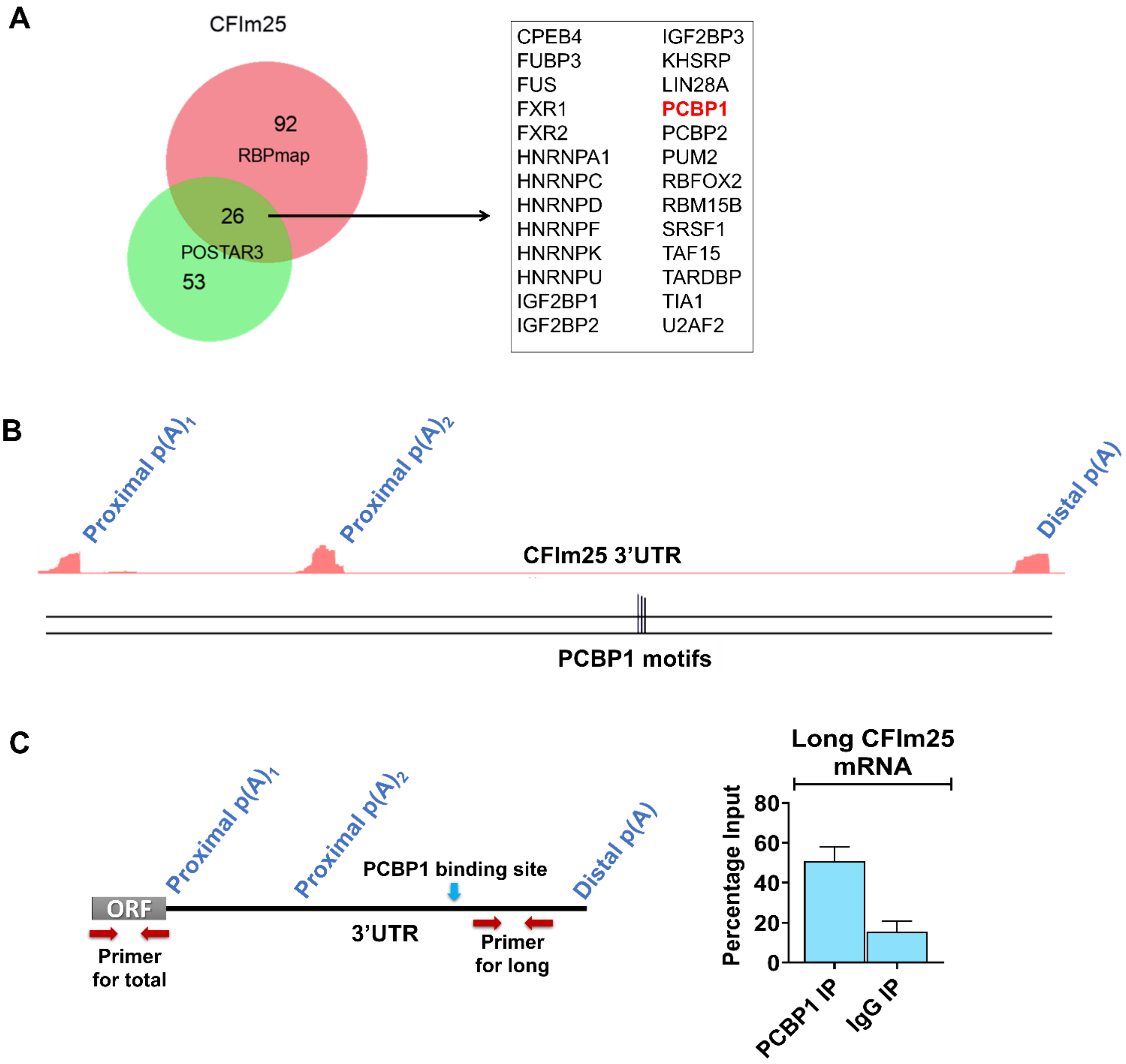


**Supplementary Fig 2. CFIm25 3'UTR RBP binding predictions.** (A) Venn diagram showing the overlap between RNA-binding proteins predicted to interact with the CFIm25 3'UTR by RBPmap (92 predictions) and POSTAR3 (53 predictions). The 26 RBPs identified by both tools are listed to the right, with PCBP1 highlighted in red. (B) UCSC Genome Browser visualization of predicted binding site for PCBP1 with the corresponding binding motifs. (C) (*Left*) Schematic showing CFIm25 mRNA structure with alternative polyadenylation sites and primer locations. (*Right*) RNA immunoprecipitation was performed using PCBP1 antibody or control IgG, followed by qPCR analysis with primers specific for the long 3'UTR isoform of CFIm25. Data represents the percentage of input RNA recovered in immunoprecipitation. Error bars represent mean ± SD (n=3). *$p < 0.05$ by Student's t-test comparing PCBP1 IP to IgG control.

**A CD38 levels by flow: representative raw data**

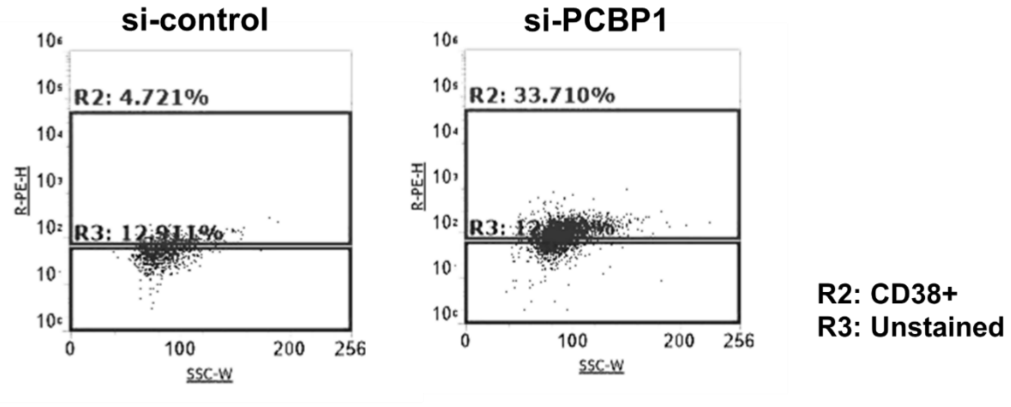


**B CD11b levels by flow: representative raw data**

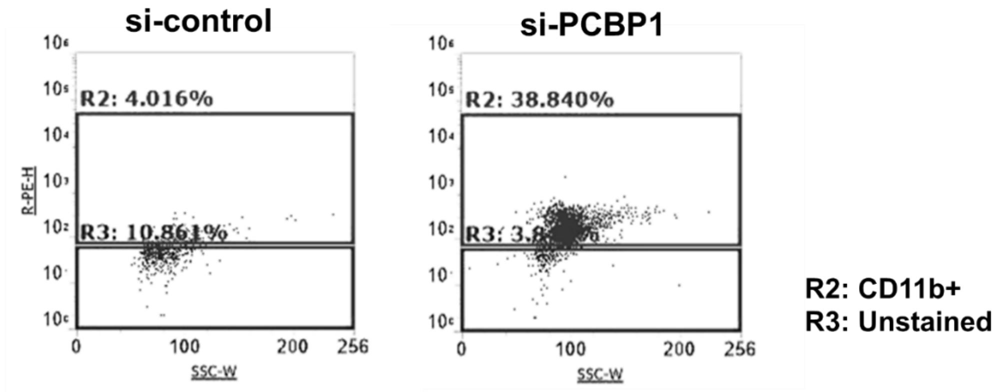


**Supplementary Fig 3. Flow cytometry raw blots (A-B**). Representative dot plots for flow cytometry of macrophage marker CD38 (A) and CD11b (B) expression (R2 gate) in undifferentiated THP-1 cells transfected with control siRNA (left) or PCBP1 siRNA (right). The percentage represents cells positive for CD38 or CD11b. Samples were prepared according to Materials and Methods and acquired on an Attune flow cytometer (Thermo Fisher Scientific), and data were analyzed using FACS diva. For analysis, samples were gated on light scattering properties to exclude dead cells and debris. Unstained control samples were used to determine the level of background fluorescence (R3). Each dot plot displays SSC on the x-axis and PE-tagged fluorochrome on the y-axis.
